# Hogs sleep like logs: wild boars reduce the risk of anthropic disturbance by adjusting where they rest

**DOI:** 10.1101/2023.03.02.530755

**Authors:** Gustave Fradin, Simon Chamaillé-Jammes

## Abstract

Many animals living in anthropized landscapes try to avoid encountering people by being active at night. By doing so, however, they risk being disturbed while at rest during the day. To mitigate this risk, diurnally resting species may be highly selective about where they rest. Here, we used GPS and activity sensors to study how wild boars (*Sus scrofa*) adjust their resting site selection and visitation patterns to the risk of disturbance by people. The data was complemented with audio recordings from animal-borne loggers to evaluate the efficacy of wild boars’ resting strategy in reducing the probability of encountering people while at rest. Generally, we found that wild boars did not specifically avoid resting near villages or roads, i.e. where the risk of encountering people is higher, if they could find sites with suitable vegetation cover. Wild boars could actually rest close to villages with very little risk of being disturbed. Resting sites located close to villages were visited more repeatedly that those located further away, suggesting that focusing on a few familiar and quiet resting sites was a successful strategy for resting undisturbed in an anthropized landscape.

## 1 Introduction

Resting is a crucial daily requirement for all animal species. It serves various purposes such as energy conservation (Riede, van der Vinne & Hut 2017; Glass, Breed, Robards, Williams & Kielland 2021), thermoregulation (Lutermann, Verburgt & Rendigs 2010), and predator avoidance (Lima, Rattenborg, Lesku & Amlaner 2005). Additionally, rest encompasses sleep, which is essential for neuro-physiological homeostasis (Schmidt 2014). The timing, duration, and location of resting is largely influenced by life-history traits such as body size and diet (Siegel 2009). As a result of physiological and ecological constraints, animals allocate a significant proportion - often more than half - of their daily time budget to resting, which takes place at specific resting sites (Campbell & Tobler 1984). These sites may vary in quality, and in particular may offer different levels of safety (Markham, Alberts & Altmann 2016; Burger, Fennessy, Fennessy & Dierkes 2020). Thus, resting site selection can be fine-tuned to minimize the risk of predation or disturbance, and the need for daily rest is likely to have an important influence on how animals use the landscape.

In many environments, animals are facing increasing anthropic pressures that can impact their resting behavior (Gaynor, Hojnowski, Carter & Brashares 2018; Wilson et al. 2020). It is well documented that many species modify their activity patterns in response to human pressures, and tend towards nocturnality to avoid being active during the day, when the intensity of human activity is highest (Gaynor et al. 2018). Such a response to anthropisation, however, exposes resting behavior to higher risks of disturbances by people. When at rest, animals may be subjected to both targeted (e.g. hunting) and untargeted (e.g. outdoor sports) disturbances, and in response should have developed strategies to reduce this risk. In particular, the spatial organization of resting may be adjusted to minimize the risk of being disturbed. Some species have been shown to fine-tune their selection of resting sites to the perceived level of risk of anthropic disturbances. For instance, wolves have been demonstrated to select more concealed resting sites after an encounter with people (Wam, Eldegard & Hjeljord 2012), and to proactively choose to rest farther from human infrastructures (Bojarska et al. 2021). Elephants have been observed returning more often to preferred resting sites, when ranging outside protected areas (Wittemyer, Keating, Vollrath & Douglas-Hamilton 2017). However, despite a few documented examples such as these, our overall understanding of adjustments of resting site selection and visitation rate to the risk of disturbance by people remains limited. Additionally, we are not aware of any study having quantified the actual success of such adjustments in reducing the likelihood of being disturbed. Generally, there is very little information about how frequently animals are disturbed while resting in the wild.

One species that offers great opportunities to learn more about these adjustments is the wild boar (*Sus scrofa*). It is a widespread species that can be found to survive and exploit even the most anthropized landscapes (Stillfried et al. 2017). When living near people, wild boars could naturally be subjected to non-targeted disturbances (e.g. by walkers and dog being walked). Additionally, as a commonly favored game species, wild boars are subject to hunting. Wild boars are known for their great behavioral flexibility and are able to adjust their patterns of activity to various factors including the season, the food availability (Keuling, Stier & Roth 2008; Brivio et al. 2017), but also the risk of human encounter (Ohashi et al. 2014; Johann, Handschuh, Linderoth, Dormann & Arnold 2020; Rosalino et al. 2022). Although they can be readily active during the day where human densities are low (Podgórski et al. 2013), they become more nocturnal when they live near people (Podgórski et al. 2013; Ikeda, Kuninaga, Suzuki, Ikushima & Suzuki 2019), and when they are hunted (Keuling, Stier & Roth 2008; Johann et al. 2020). As most outdoors activities - including recreational hunting - occur during the day, animal behaviors that take place during daytime are most at risk of disruption by people. By switching to a predominantly nocturnal pattern of activity in human-dominated landscapes, wild boars chose to expose their resting phase to a higher risk of anthropic disturbance. This likely is a successful strategy for risk mitigation, as resting is a well-known and efficient anti-predator strategy (Lima et al. 2005). However, this also means that the daily choice of a resting site is critical for minimizing the risk of encountering people. When nocturnal, wild boar are monophasic sleepers, and usually remain at rest for the entire daytime period. This implies that the choice of a specific resting site, made in the early morning, determines the exposure of the animal to the risk of disturbance for the rest of the day. The decision-making process for choosing a resting site has several components, offering several opportunities to adjust for the risks imposed by people. For example, wild boars could adjust where they rest in the landscape (e.g. distance to infrastructures, habitat types), and how they use their network of potential resting sites, coming back to some more often. It is well known that anthropic pressures can shape the spatial behavior of wild boars (Tolon et al. 2009; Stillfried et al. 2017), and several studies have investigated how their use of resting sites varies with the risk of being hunted (Maillard & Fournier 1995; Sodeikat & Pohlmeyer 2007; Scillitani, Monaco & Toso 2010; Saïd, Tolon, Brandt & Baubet 2012). However, studies rarely considered the risk of disturbances unrelated to hunting, despite their potential importance in anthropized landscapes (Marzano & Dandy 2012).

Here, we present a study of the resting strategy of wild boars living in a rural Mediterranean landscape in the South of France. Using activity and GPS data collected on collared animals, we examined how wild boars modify their resting site selection in response to the risk of anthropic disturbance. This includes the assessment of (i) the potential avoidance of areas where meeting people is more likely (i.e. close to villages and roads), (ii) the selective re-use of previously visited sites, depending on their location in the landscape with regards to the risk of being disturbed. We also estimated, across the landscape, how frequently wild boars left their resting site and engaged in a significant and potentially costly relocation to a secondary site. We also used audio recordings from animal-borne loggers to try to detect human activities interfering with wild boars’ rest. Overall, our study provides insights into the effectiveness of behavioral adjustments in reducing the risk of disturbance of rest by animals living in anthropized landscapes.

## 2 Materials & methods

### 2.1 Study Sites

We studied wild boars in two areas in southern France: the *Gorges du Gardon* area (43.93°N; 4.38°E) and the *Pic Saint-Loup* area (43.74°N; 3.88°E). Both sites are very similar. Altitude is ∼100m, and the landscape is a mosaic of agricultural lands, densely vegetated patches (shrublands and forests), and villages (covering respectively 38%, 48%, and 11% of the land in the *Gorges du Gardon* area, and 55%, 29%, and 12% in the *Pic St-Loup* area). Vineyards make up most of the agricultural lands. The natural vegetation patches are mostly Mediterranean *garrigues*, with thick understory and dominated by dense evergreen oak (*Quercus ilex*) or Aleppo pine (*Pinus halepensis*) tree stands. Villages have hundreds to a few thousand inhabitants, and the general population density is approximately 280 inhabitants/km², with some isolated houses also spread throughout the landscape. Both sites are located about ten kilometers away from a large city (Nîmes for the *Gorges du Gardon* area, Montpellier for the *Pic Saint-Loup* area). Wild boars are mostly hunted in drive hunts involving an average of twenty hunters (usually between ten and forty) and about three to fifteen hounds. The open season for drive hunt lasts from the 15^th^ of August to the end of February or March, depending on the year and the study area. In addition to the risk induced by hunting, wild boars could potentially be disturbed, at both study sites, by agricultural activities, and by recreational outdoors activities (e.g. biking, hiking, dog-walking, etc.). The lack of natural predators in the study areas guarantees that human activity represents the only potential risk of disturbance for resting wild boars. Part of the *Gorges du Gardon* area is a densely vegetated military zone where hunting is allowed, but where the other human activities differed from elsewhere, although we could not evaluate how it could affect wild boars’ behavior.

### 2.2 Captures and tracking

From 2018 to 2022, we captured wild boars with corn-baited traps, using a protocol approved by the ethical committee of the French Ministry of Research (APAFIS#20279-2019041522576537v3). We equipped 22 animals weighting more than 40kg with a GPS collar, using either anesthesia or a small immobilization box. The collars acquired a GPS location every 30 minutes, and a measure of activity level (hereafter ACT) every 5 minutes. ACT was calculated as the average of 8Hz acceleration measurements over three orthogonal axes, and was available to us on a dimensionless scale ranging from zero (indicating no movement of the collar over the 5-minute interval) to 255. As values were highly correlated between axes, we only used the ACT data corresponding to the anteroposterior axis. We recovered nine collars through the activation of their remote drop-off system, two after the animals had been killed by a collision with a vehicle, six after the animals had been shot by local hunters, and five after they had felt off the animals prematurely. Out of these five collars, three dropped less than fifteen days after deployment and we excluded the corresponding animals from the study. The average survey period was 126 days long (SD=53), with the first three days after capture excluded from the analysis to eliminate any abnormal behavior. The activity profiles of the animals clearly showed that most were active at night (example in Figure 1a), with one phase of inactivity spanning the daylight hours. Only two animals were significantly active during the day during parts of their survey period and were excluded from the analysis to focus only on daytime resting strategies. Seventeen animals (twelve males and five females) remained in our final dataset. Two males and two females had been using extensively the military zone. No pair of collared animals moved together as part of the same sounder. Due to the very limited female data, we excluded them from the analyses and only present results from the analysis of male data. Note however that exploration of the female data does not suggest that their behavior differ largely from that of males, with regard to aspects investigated here (not shown). Also, for brevity, we hereafter refer to the animals studied in the analyses as ‘wild boars’, without specifying their sex each time.

**Figure 1.**
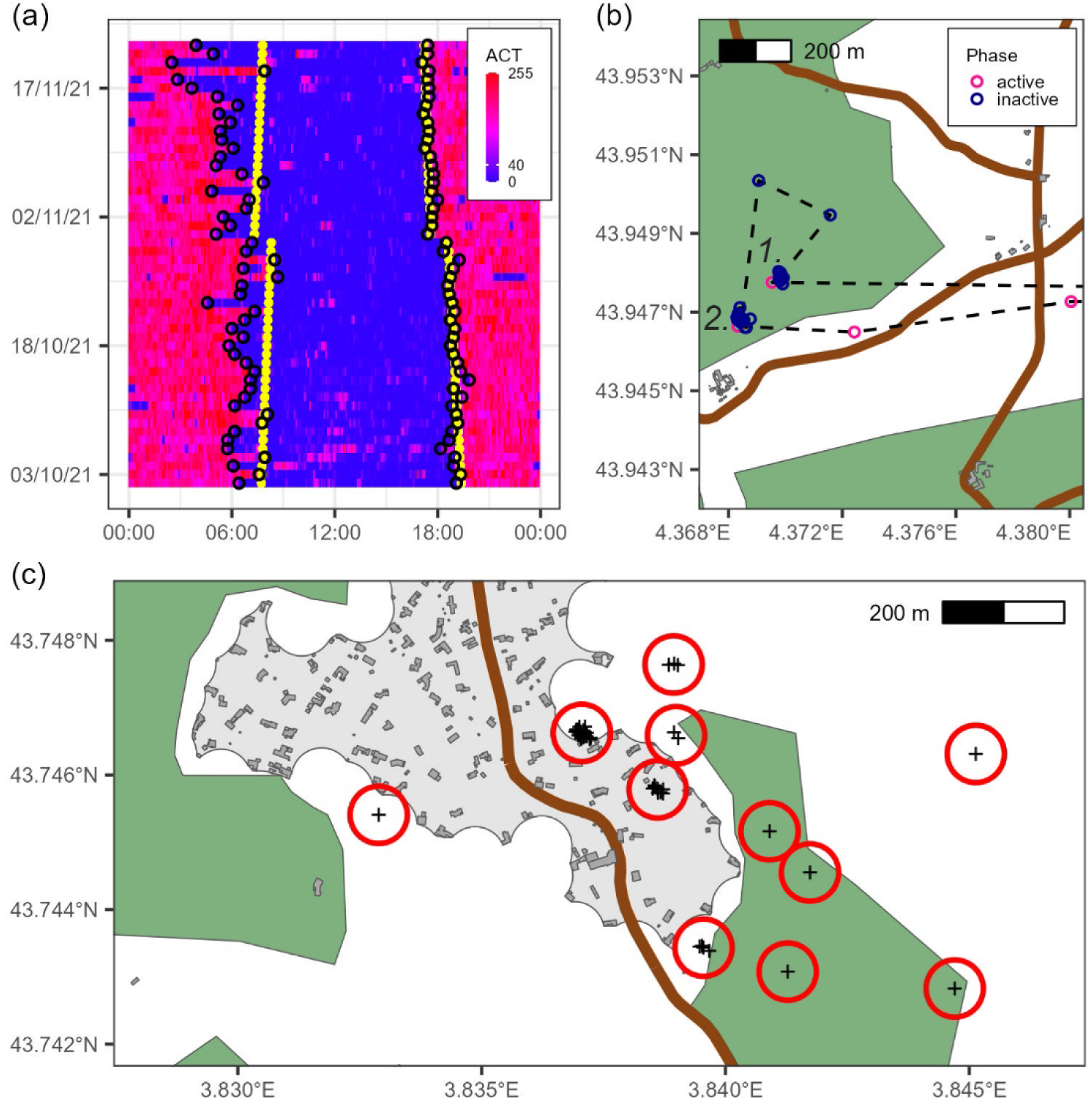
(a) Example of the activity pattern of a wild boar. The color codes for the value of the ACT variable (blue=low activity, red=high activity). The black circles indicate the start and end times of the inactive phases. The yellow dots indicate the times of sunrises and sunsets. (b) Example of a wild boar’s relocation during an inactive phase. The black dashed line represents the trajectory linking GPS fixes (pink and blue circles). The animal first rested in (1) and then relocated to spend the rest of the inactive phase in (2). (c) Example of the resting site revisitation pattern of a wild boar. The daily resting locations (black crosses) are clustered into resting sites (red circles). In (b) and (c), buildings are represented in dark grey, villages in light grey, roads in brown, and densely vegetated patches in dark green.

### 2.3 Data preparation

#### 2.3.1 When did resting occur?

To study where wild boars rested, we first needed to determine when they rested each day. Upon visually examining their activity patterns (Figure 1a), it was evident that wild boars consistently divided their 24-hour cycle between an inactive phase (primarily resting with some occasional activity) and an active phase (during which they could occasionally rest). To recover the start and end times of these inactive phases, we first classified each 5-minute period as ‘resting’ (ACT≤40) or ‘active’ (ACT>40). Followed by a few other steps, detailed in Appendix 1, this allowed us to identify a continuous inactive phase spanning the daytime hours each day, separated from the previous and next inactive phases by an active phase. During this process, we disregarded the short bouts of activity that occurred between long-lasting ‘resting’ bouts, which were likely due to slight movements during resting. The resulting patterns closely matched those anticipated based on visual inspection (see Appendix 1).

#### 2.3.2 Where did resting occur?

We used the GPS data to determine where the wild boars rested during each inactive phase. Usually, the entire inactive phase was spent at a single location. On some occasions, however, the wild boars would start resting in one place and then move to another location to spend the rest of the inactive phase. We called such events ‘relocations’ (example in Figure 1b). To identify relocations, we looked for any substantial movements recorded in the ACT data during the inactive phases. We compared the mean GPS position of the animal before and after the high ACT values, and if those locations were more than 100 meters apart, we considered that the animal had relocated. See Appendix 1 for more details.

Throughout their lives, wild boars tend to return to rest at sites where they have rested before. To test whether this could be affected by where the resting sites are in the landscape, we first had to define ‘resting sites’ (hereafter called ‘RS’, example in Figure 1c). We did so by using one location per inactive phase and performing a hierarchical clustering analysis, using an average linkage, to define clusters that were more than 50 meters apart. All the locations that fell within the same cluster were considered to be revisitations of the same RS. To ensure that the positions of the RSs reflected unconstrained choices made by the wild boars when selecting a resting place, we ignored all places visited after a relocation, as their position was constrained by the context of the relocation (the previous resting location and the reason for the relocation).

#### 2.3.3 Environmental variables

We first characterized the vegetation cover at each RS. Our study sites could be coarsely divided into agricultural lands, densely vegetated areas, and villages. Wild boars rarely rested inside villages, and when they did, they always stayed at the outskirts, in the transitional zones between villages and agricultural lands. To simplify, we characterized vegetation cover as a binary variable, indicating whether the RS was located within a densely vegetated patch or not. We used the CORINE Land Cover 2018 data to differentiate between large patches of dense vegetation and agricultural areas with scattered small thickets and hedgerows. See Appendix 2 for more details.

Then, we used OpenStreetMap (OSM) to calculate the distance of each RS to two features associated with higher risks of human disturbance: villages and roads. We laid out the road network by combining the ‘motorway’, ‘trunk’, ‘primary’, ‘secondary’, and ‘tertiary’ road categories from OSM. This way we excluded the residential roads, the roads with very low traffic, and the forest tracks. We also used OSM data to determine the edges of villages. We did so by applying a buffer of 50 meters around OSM’s polygonal layer for buildings and then shrinking the resulting polygons with a buffer of -50 meters. This allowed us to identify villages as clusters of buildings closer together than 100 meters, while aligning their edges with the peripheral buildings’ edges. Any such polygon larger than 1.5 hectares was considered a village and visual inspection confirmed the adequacy of this approach. We calculated the distances of each RS to the nearest road, and village edge (with negative distances indicating locations inside villages).

Finally, we associated each visit of a RS to the status of the hunting season the day it occurred. Drive hunting is the primary hunting technique for wild boars in the study areas, and the only one that targets resting animals. Therefore, we defined the hunting season (HS) based on the dates of the open season for drive hunting. Drive hunts were only permitted on ‘hunting days’ (Wednesdays, Saturdays, Sundays, and bank holidays). During the non-hunting season (NHS), drive hunts were prohibited, except with a specific authorization from the local authority, concerning a negligible number of hunts. We did not account for the Covid-19 lockdowns because they never affected the hunting patterns during the periods in which we monitored the animals. Although lockdowns may have reduced the probability of untargeted disturbances, in our study this only affected 28 days for four animals.

### 2.4 Data analysis

#### 2.4.1 Habitat selection for resting sites

We investigated how the selection of RS by wild boars varied with vegetation cover and proximity to villages and roads. We estimated this selection within the home ranges of the animals, defining the home range as the contour of the 90% utilization distribution, estimated with a standard kernel density approach, from GPS locations collected during the active phase. We used the selection ratio as a metric of habitat selection. A selection ratio above one corresponds to selection, and a value below one corresponds to avoidance. We followed Chamaillé-Jammes (Chamaillé-Jammes 2020) who showed how the selection ratio can be computed along continuous variables from a standard resource selection function (RSF). For the RSF, the available RS were created by drawing, within the home range of each wild boar, a thousand random points per actual RS. These available RS were characterized using the environmental variables described above. We then estimated how the selection ratio for RSs varied with whether they were located inside or outside densely vegetated patches, and with their distance to roads and villages. We did so by fitting the RSF using a generalized linear mixed model (GLMM), using the type of location (used vs. available) as response variable and a binomial distribution for errors. Predictor variables were the vegetation cover (inside or outside a densely vegetated patch) in interaction with the distances to roads, and to villages. We also included individual animal identities nested within study sites as random intercepts. We also expected the hunting season to influence how wild boars selected RSs. However, many RSs had been visited during both during the HS and the NHS, so that we could not simply include the hunting season in the RSF as a predictor variable. We thus decided to run two separate models: one with the RSs visited at least once during the HS, and one with the RSs visited at least once during the NHS.

#### 2.4.2 Resting sites’ revisitation rate

Some RSs were preferred and visited more often than others. We investigated how the proximity of villages and roads influenced the frequency of visitations of RSs. We did so by fitting a GLMM on the number of visits to each RS, using a zero-truncated Poisson regression. As predictor variables, we used the vegetation cover (inside or outside densely vegetated patches) in interaction with the distances to roads, and to villages. The length of the survey period would obviously influence the number of times a RS could be visited, and we therefore included this variable (log-transformed to account for the link function of the Poisson regression) as an offset in the model. Finally, we included individual animal identities, nested within study sites, as random intercepts. We did not account for the status of the HS, as any many RS were visited both during and out of the HS.

#### 2.4.3 Relocation events

We used relocation events to gain insight into how often wild boars are disturbed while at rest. We modeled the probability of a wild boar to relocate as a function of the distances to roads and to villages, in interaction with whether the initial RS was inside or outside a densely vegetated patch. A fixed effect variable to account for the risk of being hunted was also included. In the analysis of habitat selection of RSs above, we implicitly studied proactive adjustments of resting behavior to human activity and assumed that wild boars could behave differently during the HS (vs. during the NHS). In contrast, here, we expected that relocation events could be influenced not only by the HS being open or closed, but also by wild boars being actually searched for and chased by dogs, i.e. by whether the day was actually a hunting day or not. Therefore, we used a 3-category variable coding for ‘HS and hunting day’, ‘HS and non-hunting day’ and ‘NHS’. We also included individual animal identities, nested within study sites, as random intercepts. We also modeled the distance traveled between RSs each time a relocation event occurred using a similar model and the same predictor variables.

#### 2.4.4 Audio data extraction and analysis

Six of the collars, deployed on three females and three males, had an audiologger (Latorre, Miquel & Chamaillé-Jammes 2021), i.e. an autonomous microphone, which recorded an average of 16 days (SD=9) of continuous audio data after capture. We used these data to gain insight into what exactly happened right before relocations occurred. Nine relocation events occurred during a period with audio monitoring, with an average distance traveled of 170 meters (maximum=317m). For each relocation event, we precisely identified the moment the wild boar stopped resting (a sharp rise in ACT) and extracted the audio from 40 minutes prior to this moment to 5 minutes after. This way, we ensured that the extract started while the wild boar was resting and quiet, and ended after it had started to relocate. For each of those ‘relocation’ audio tracks, we also extracted one ‘control’ track, recorded at the same time of day but on a different date. Tracks are available in Appendix 3. We randomly sorted all tracks, listened to them while viewing the corresponding spectrogram, and noted any sounds that could be identified. In particular, we focused on sounds related to human activities (e.g. motor sounds, dog barks, human voices, gunshots) that could interrupt the resting of wild boars. Of course, this approach might have missed some disturbances if they occurred silently or if the wild boars were able to detect danger and relocate before any sound could be picked up by the microphone. However, a sensitivity trial we conducted under realistic conditions suggested that the microphones were sensitive to even faint distant sounds: we placed a microphone in the vegetation, facing the ground, and spoke in a calm voice several meters away from it. We were able to identify normal speech both in the audio recording and in the spectrogram when the experimenter stood as far as 30m away from the microphone, facing it (20m when facing the opposite direction).

## 3 Results

### 3.1 Timing and duration of inactive phases

The wild boars studied spent most of their time at rest. On average, 58.5% (SD=4.4) of their time budgets corresponded to the inactive phase, which typically started at the last hours of the night, and spanned throughout daytime. Activity patterns were mostly synchronized with sunset times throughout the year (examples in Figure 1a and in Appendix 1). The end of the inactive phase occurred within 30 minutes around sunset in 44% of cases (and within 90 minutes in 80% of cases), whereas it started within 30 minutes around sunrise in only 24% of cases (and within 90 minutes in only 53% of cases).

### 3.2 Patterns of resting site selection

#### 3.2.1 Habitat selection for resting sites

As expected, wild boars favored resting in dense vegetation patches and generally avoided resting at sites that were in more open habitats (Figure 2; Table 1). During the NHS, the proximity of roads or villages had no effect on the selection of sites by wild boars, as long as they were in dense vegetation (Figure 2a; Table 1). Sites in more open vegetation were avoided when they were close to roads, but this avoidance disappeared, maybe shifting to a selection, when they were far from roads (Figure 2a; Table 1). Note however the large confidence intervals, caused by the fact that little data was available for those sites, that are rare in the landscape. During the HS, wild boars also did not adjust their selection to the distance to the closest village (Figure 2b; Table 1). In contrast to the NHS, they however did select more strongly sites located close to roads when they were in dense vegetation (Figure 2b; Table 1). The proximity of roads had no effect on the selection of sites in more open vegetation, and they were always avoided in this season (Figure 2b; Table 1).

**Figure 2.**
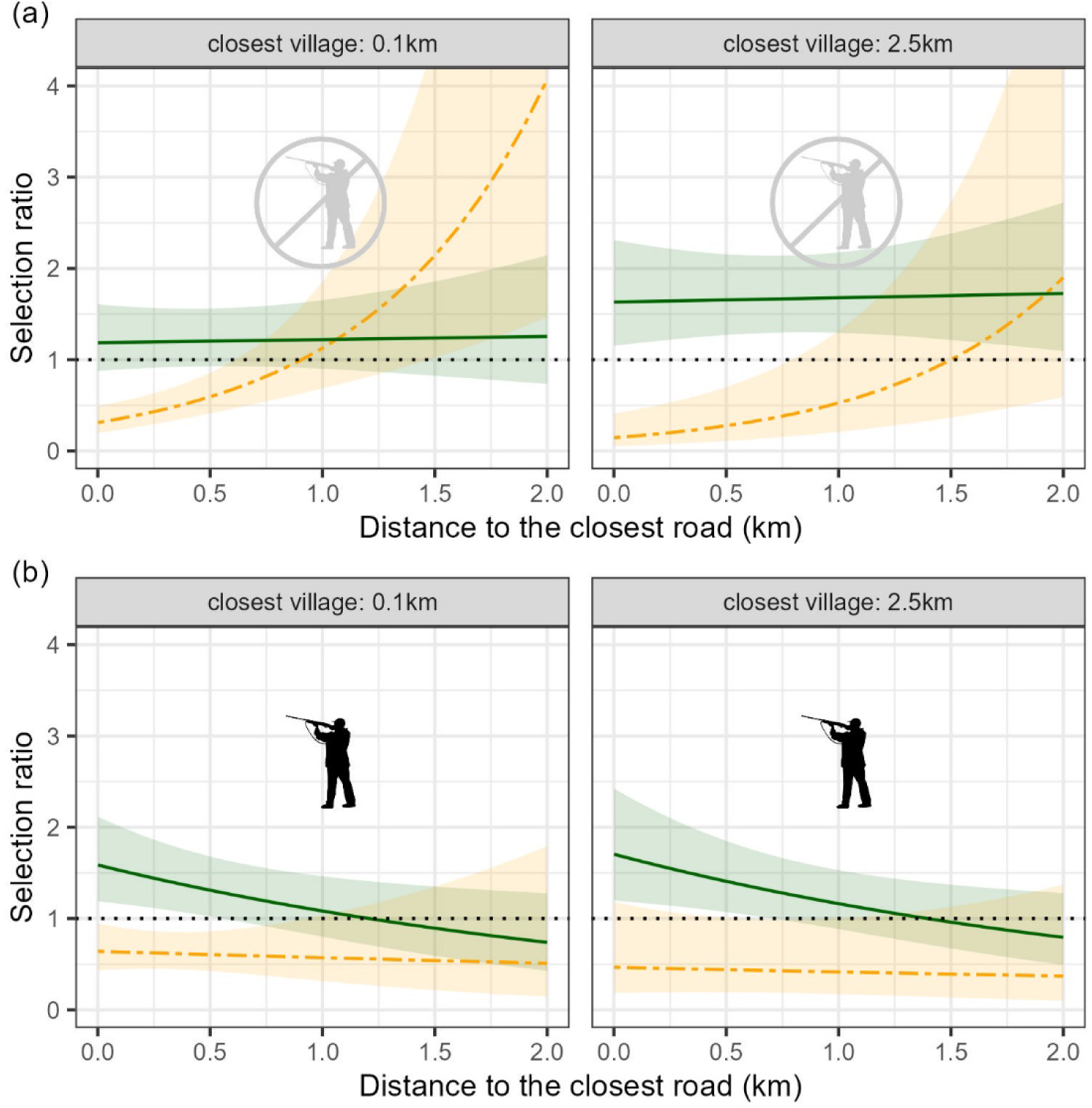
Estimation of the selection ratio for resting sites by wild boar during the non-hunting season (a) and the hunting season (b), for different distances to roads and villages. Selection for densely vegetated patches is represented in a dark green, solid line. Selection for non-densely vegetated patches is represented in a yellow, dashed line. A selection ratio above one indicates that sites are selected, and a selection ratio below one indicates that sites are avoided. Lightly-colored ribbons show the 95% confidence intervals.

**Table 1.** Statistics of the models estimating the selection ratio for resting sites.

| Variable | $\beta$ | SE | z-value | p-value |
| --- | --- | --- | --- | --- |
| **Model: Hunting season** |  |  |  |  |
| Intercept | -7.34 | 0.21 | -35.44 | <0.001 |
| inside densely vegetated patches | 0.89 | 0.26 | 3.46 | <0.001 |
| d(road) | -0.12 | 0.37 | -0.31 | 0.75 |
| d(village) | -0.13 | 0.21 | -0.62 | 0.53 |
| inside densely vegetated patches x d(road) | -0.27 | 0.40 | -0.67 | 0.5 |
| inside densely vegetated patches x d(village) | 0.16 | 0.23 | 0.70 | 0.48 |
| **Model: Non-hunting season** |  |  |  |  |
| Intercept | -8.04 | 0.24 | -33.20 | <0.001 |
| inside densely vegetated patches | 1.29 | 0.29 | 4.45 | <0.001 |
| d(road) | 1.28 | 0.31 | 4.10 | <0.001 |
| d(village) | -0.32 | 0.24 | -1.34 | 0.18 |
| inside densely vegetated patches x d(road) | -1.26 | 0.35 | -3.58 | <0.001 |
| inside densely vegetated patches x d(village) | 0.45 | 0.25 | 1.77 | 0.08 |

'd(village/road)' stands for 'distance to the closest village/road'.

#### 3.2.2 Resting sites’ revisitation rate

Wild boars regularly revisited RSs (51% were visited more than once), and some were visited more often than others (Figure 1c). RSs located close to villages were visited more often than those located further away, irrespectively of whether they were in dense vegetation or not (Figure 3a; Table 2). Although the visual inspection of the model predictions suggested a positive relationship between the frequency of visits to RSs and their distance to the closest road (Figure 3b), the relationship was not statistically significant, irrespectively of whether they were in dense vegetation or not (Table 2).

**Figure 3.**
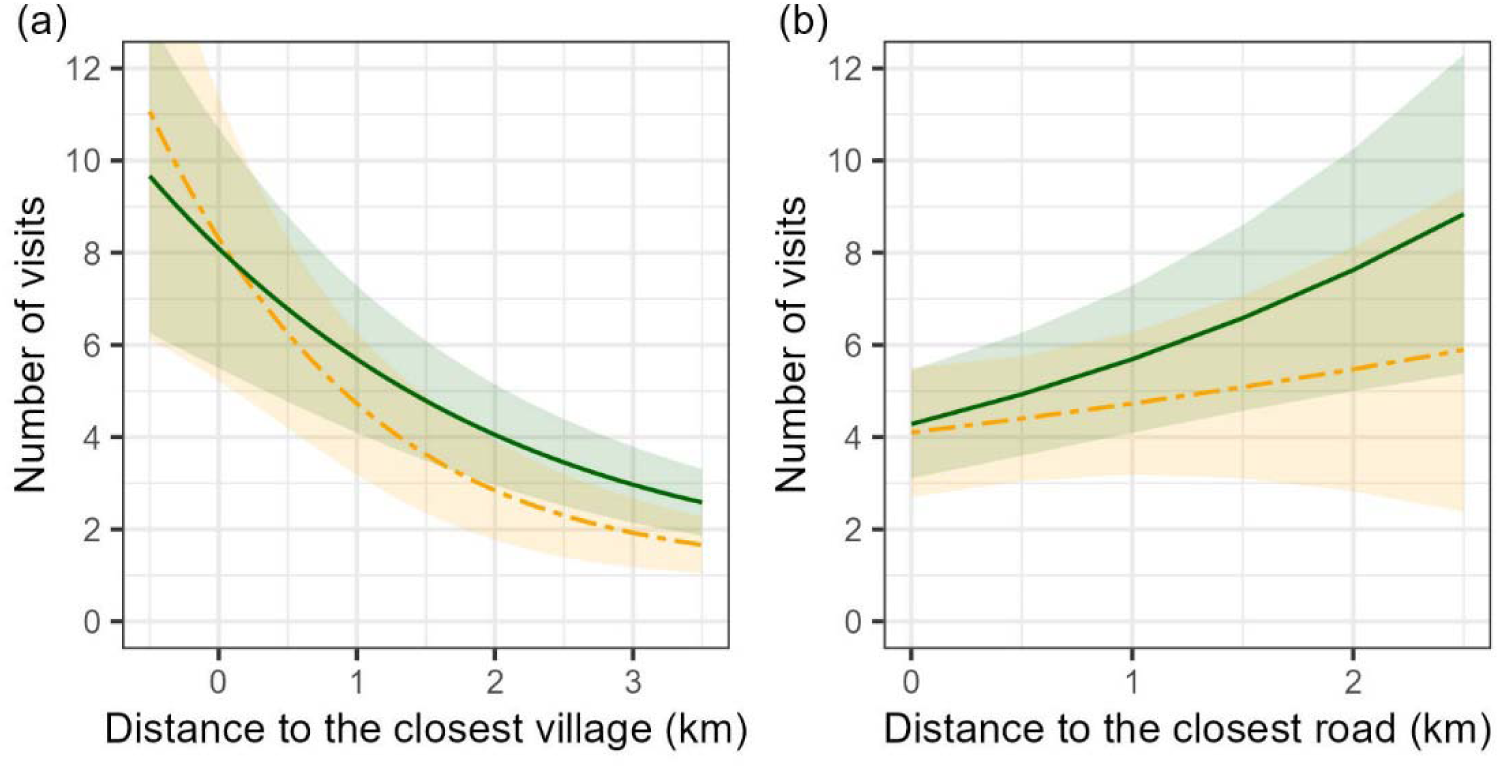
Estimation of the number of visits to resting sites (over a 6-month period), according to the distance to the closest village (a) and to the distance to the closest road (b). The estimation for the resting sites located inside densely vegetated patches is represented in a dark green, solid line. The estimation for the resting sites located outside those patches is represented in a yellow, dashed line. Lightly-colored ribbons show the 95% confidence intervals.

**Table 2.** Statistics of the model estimating the number of visits to resting sites.

| Variable | $\beta$ | SE | z-value | p-value |
| --- | --- | --- | --- | --- |
| **Model: Number of visits** |  |  |  |  |
| Intercept | -3.23 | 0.17 | -19.29 | <0.001 |
| d(village) | -0.57 | 0.13 | -4.36 | <0.001 |
| d(road) | 0.15 | 0.14 | 1.08 | 0.28 |
| inside densely vegetated patches | -0.17 | 0.11 | -1.57 | 0.12 |
| inside densely vegetated patches x d(village) | 0.22 | 0.14 | 1.54 | 0.12 |
| inside densely vegetated patches x d(road) | 0.14 | 0.15 | 0.94 | 0.35 |

'd(village/road)' stands for 'distance to the closest village/road'.

### 3.3 Relocations

The probability that a relocation occurred during the inactive phase was very low during the NHS (0.08, 95%CI [0.05-0.11]; Figure 4a; Table 3). In addition, when a relocation happened during the NHS, the distance traveled was virtually always less than 500 meters (Figure 4b). When the animal was initially resting in dense vegetation patches, neither the distance to the closest village, nor the distance to the closest road affected the probability of a relocation (Figure 4c; Table 3), or the distance traveled on relocations when they occurred (Table 3). When the animal was initially resting outside dense vegetation patches, there were a few significant effects of distance to the closest village or road on the probability of relocation and on the distance traveled, that were likely not biologically relevant (small effect or large confidence interval) and possibly caused by the low sample size in this condition (Table 3). Although the probability of relocation remained relatively low in the HS, it did increase two-fold during the hunting days of the HS (0.17, 95%CI [0.11-0.27]; Figure 4a; Table 3). Also, almost all of the few long-distance relocations (17 out of 21 of the relocations >500 meters) happened during the HS (Figure 4b).

**Figure 4.**
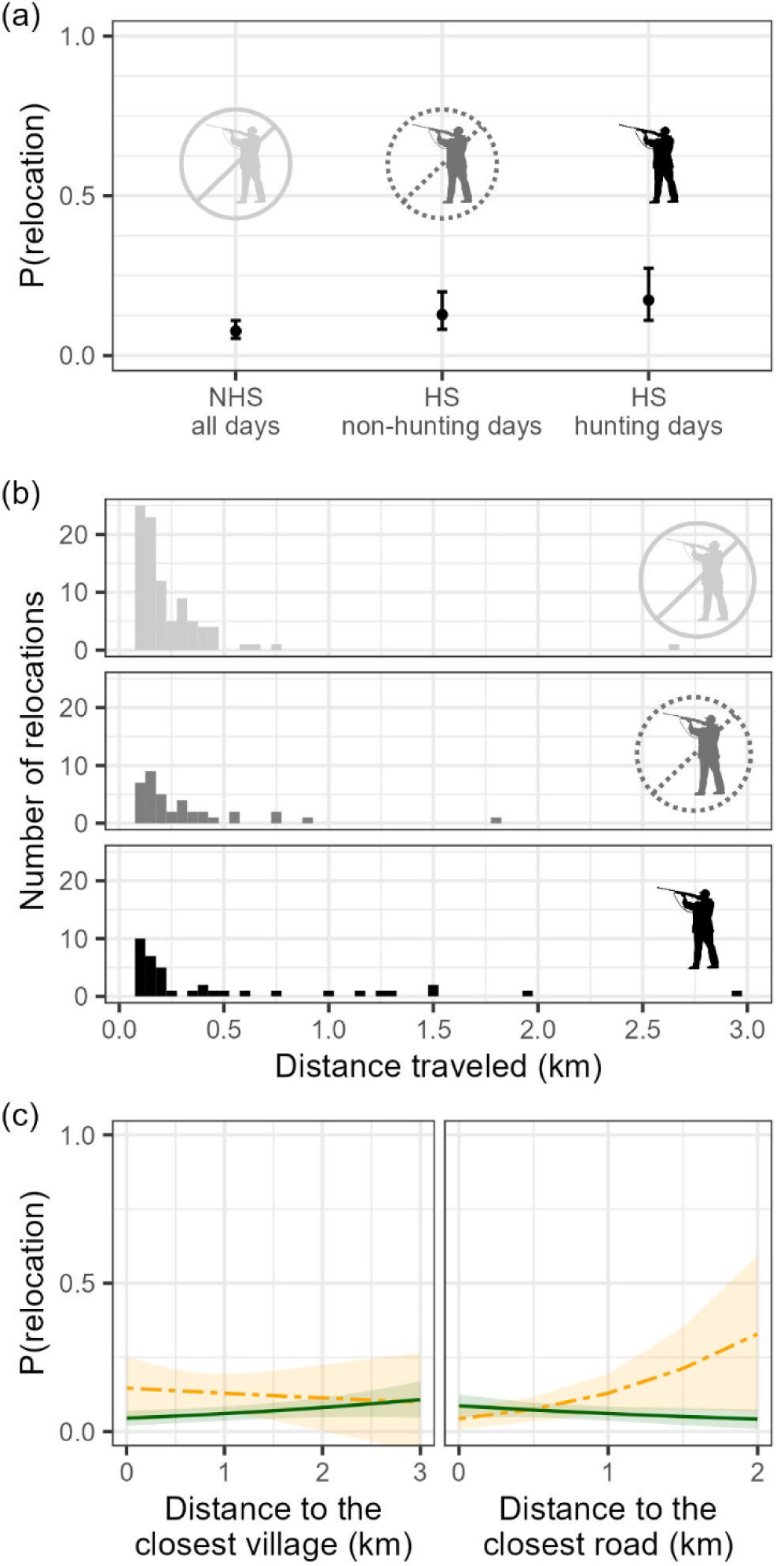
(a) Estimation of the probability of relocation during the hunting season and hunting days (Wednesdays, Saturdays, Sundays, and bank holidays), during the hunting season and non-hunting days, and during the non-hunting season. (b) Distribution of the distances traveled during relocations, according to hunting periods (same categories as in (a)). (c) Estimation of the probability of a relocation, during the non-hunting season, according to the distance to the closest village and to the closest road. The estimation for the resting sites located inside densely vegetated patches is represented in a dark green, solid line. The estimation for the resting sites located outside those patches is represented in a yellow, dashed line. Lightly-colored ribbons show the 95% confidence intervals.

**Table 3.**
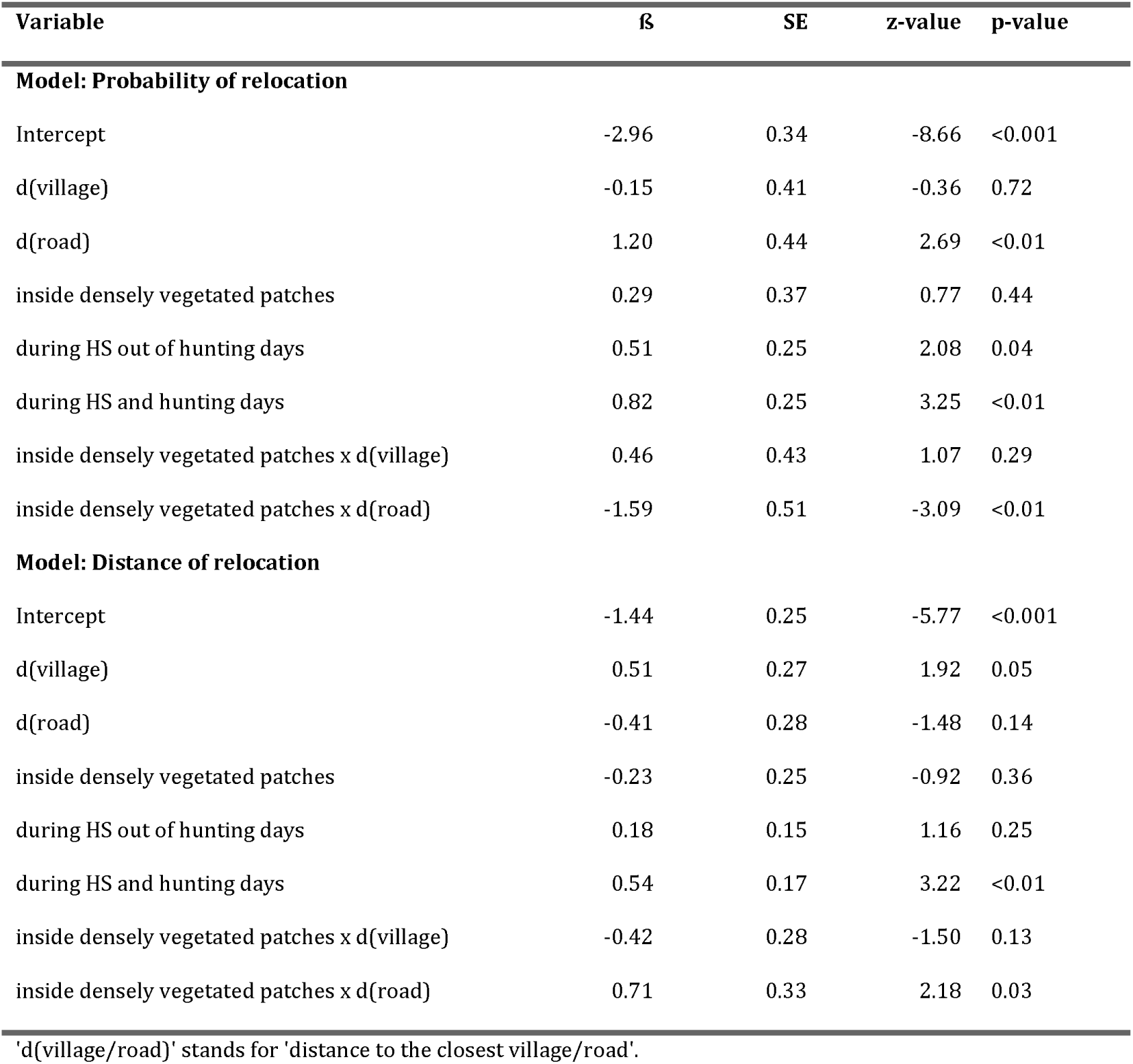
Statistics of the models estimating the probability and the distance of relocations.

In both the audio tracks extracted from before relocations and the control tracks, the wild boars could be heard resting quietly, usually snoring, breathing calmly and grooming. Although we could hear vehicles passing at close distance in two control tracks, the wild boars did not seem to react to those sounds. In 8 extracts corresponding to a relocation, we could hear the wild boar walking away, without any disturbance being identifiable in the audio. In the ninth extract, which was recorded on a Sunday during the HS, we could clearly identify a hunting event by the barks of a pack of dogs and a gunshot, right when the wild boar started to move away (Appendix 3). We did not hear any other potential disturbance linked to human activity in the selected extracts.

## 4 Discussion

Resting is an important part of an animal’s life, both in duration and function. Yet, our knowledge of the flexibility of animals’ resting behavior remains limited. In particular, we know little about how animals adjust their resting behavior to the risk of human disturbances, how often actual disruptions of resting phases occur, and how animals respond to them. The wild boars we studied allocated more than half of their time budget to resting, mostly during daytime, and thus preferred to be exposed to anthropic disturbances when resting than when foraging. This pattern has been observed in other populations of wild boars (Brivio et al. 2017; Johann et al. 2020; Rosalino et al. 2022) but also in a variety of other species (Gaynor et al. 2018) living in anthropized landscapes. We questioned whether wild boars tried to minimize the risk of being disturbed through resting site selection, and in such a case, whether they were successful.

As expected, wild boars always favored resting under the cover of densely vegetated patches, which is certainly the best way to reduce the risk of being disturbed. Additionally, we expected the selection of RSs to vary with the risk of encountering people. Surprisingly, however, we did not find any effect of the proximity of villages on the selection of RSs. As resting near villages is undoubtedly associated with costs and benefits, linked to both resource acquisition and safety, these must probably offset each other so that there is not net advantage of selecting to rest near or far from a village. The fact that the proximity of villages did not affect habitat selection for RSs however contrasted with the strong effect it had on the fidelity to RSs. Wild boars came back to rest at the same spot more often when it was close, or even within, a village. We tentatively suggest that predictably quiet places, suitable for resting, are rare near villages, and that wild boars rely heavily upon them. Our study shows that the need to find suitable RSs does not prevent wild boars from exploiting areas with high degrees of proximity with people, as they can compensate a general increased probability of disturbance by selecting more intensely some specific RSs that they know are safe. This plasticity in wild boars’ resting strategy is certainly critical to explain their success in subsisting, and sometimes thriving, in a wide diversity of habitats, even those heavily urbanized (Stillfried et al. 2017).

The effect of roads on wild boars’ RS selection was more complex. During the NHS, wild boars selected to rest in densely vegetated patches irrespectively of their distance to roads. In contrast, wild boars avoided resting outside densely vegetated patches only close to roads, and they selected, or at least were neutral to, such sites if they were far from roads. This could be because the risk of being disturbed was lower there. However, note that areas outside densely vegetated patches and far from roads are rare in the landscape, leading to large confidence intervals that warrants caution in interpretation. Wild boars’ response to the proximity of roads changed during the HS, as they selected densely vegetated sites more strongly when they were close to roads, and did not select sites in more open vegetation even far from roads. We argue that this could reflect a proactive strategy of wild boars to minimize hunting risk. Hunters tend to focus their effort away from main roads as they make hunting dangerous for people and dogs. Areas near roads are thus safer than elsewhere, and exposure in the open, far from roads is likely particularly unsafe for a wild boar during the HS.

Resting strategies that involve behavioral adjustments to the risk of disturbances by people, as demonstrated here for wild boars, have been documented before (Scillitani, Monaco & Toso 2010; Wittemyer et al. 2017; Bojarska et al. 2021). Previous studies however rarely evaluated how successful these adjustments were at reducing this risk. We addressed this gap by studying the probability of wild boars relocating during their resting phases, a possible indicator of disturbance, depending on the location of the RSs. Our results generally suggest that the RS selection strategy of wild boars was effective in keeping the probability of being disturbed low, even in places where chances of meeting people is high. Wild boars rested at all distances from villages, and yet the probability of them relocating during the inactive phase was just as low close to villages than several kilometers away from them. This supports our assertion that, close to villages, wild boars use particularly quiet spots where encounters with are unlikely, and re-use them over time. They could thus rest in anthropized landscape with little cost.

Of course, not all disturbances may lead to a relocation. It is well known that wild boars may hide rather than flee when hunted (Scillitani, Monaco & Toso 2010; Thurfjell, Spong & Ericsson 2013), and generally wild boars may sometimes be disturbed and not leave their RS. We actually observed this response in 62% (8 out of 13) of the experimental disturbances that we led on resting wild boars (personal observations). Although any disturbance can affect the ‘quality’ of resting (e.g. through sleep deprivation), keeping quiet in response to an untargeted disturbance is likely a good strategy, as it avoids the costs and risks associated with relocating. Besides, non-targeted disturbances are often short-lived, and will only reduce resting time marginally. Additionally, in our study, the wild boars remained inactive for 14 hours per day on average, which was longer than the total sleep time previously recorded in domestic pigs (7.8 hours) (Campbell & Tobler 1984) and in free-ranging wild boar (10.6 hours) (Mortlock et al. 2022). This suggests that disturbances that do not entice a relocation likely have only a minor implication for fitness. In some cases, wild boars might not even respond to a stimulus that we would have assumed stressful for them. For instance, upon opportunistically listening to the rest of the audio data, we heard a dog barking from what seemed like a close distance, but the wild boar did not react and continued to snore calmly throughout the event (Appendix 3). This extract supports the hypothesis that resting wild boars are little sensitive to untargeted disturbances. Given the low frequency of relocations, and the little impact expected from disturbances that do not entice relocations, it seems safe to conclude that wild boars, in our study area, are little affected by human activity during resting. Unsurprisingly though, this was less true during the hunting season as hunting is a significant source of targeted disturbance for wild boars in this landscape. The probability of relocation was highest during hunting days of the HS, and long-distance relocations (>1km) were virtually only observed during these days. These could correspond to actual hunts, with dogs chasing the animal over large distances. The moderately greater probability of relocation in non-hunting days of the HS might reflects a higher responsiveness of wild boars to even non-targeted disturbances, during the HS.

Of course, not all relocations may have been a response to a disturbance. In 9 out of 10 audio extracts collected at the onset of a relocation, the animal could only be heard resting quietly and snoring, before leaving the area for no apparent reason. It is possible that we did not hear a person or a dog that was nearby, which the wild boar would have smelt, or heard, and reacted to. It is also possible that the wild boar simply woke up and left for a new area for a different reason (e.g. weather change). If such natural daily movement occur during the inactive phase, this would strengthen our general interpretation that people rarely disturb wild boars once they have selected a RS.

In conclusion, our study shows that wild boars are efficient at exploiting areas near people, while still finding suitable sites where to rest. They do so by selecting sites with high cover, by responding to the proximity of human presence in ways that can change with the hunting season, and by revisiting preferentially the safest sites near human settlements. It is apparently a successful strategy, as they rarely have to relocate far away in response to a disturbance, apart when targeted by hunting. As more and more habitats become subject to anthropisation, the need to find suitable sites where to rest is likely to become a constraint for a variety of animal species. In such circumstances, species with a high behavioral and ecological flexibility are likely to be favored, in particular through the adjustment of their resting patterns. Although successful temporal adjustments of activity patterns in response to human pressure had been extensively described (Lowry, Lill & Wong 2013; Gaynor et al. 2018), our study shows that resting site selection is another efficient way for animals to avoid encounters with people.

## Acknowledgments

This work was supported by the REPOS project through the I-site Montpellier Université d’Excellence (grant number ANR-16-IDEX-0006) and by the François Sommer foundation. We are indebted to the various landowners who permitted to work on their land, to S. Perret, D. Degueldre, G. Fréchet, J. Nicot for assistance with wild boar captures, and to R. Mathevet and J.-M. Chanabe for some aspects of project management. D. Cornelis kindly offered the first collars used in the study. L. Latorre and J. Miquel developed the audiologgers. M. Valeix kindly reviewed a draft of the manuscript. We also thank the map data copyrighted OpenStreetMap contributors for using the map data available from https://www.openstreetmap.org.

## Author contributions

**Gustave Fradin:** Conceptualization (equal); data curation (lead); formal analysis (lead); investigation (supporting); visualization (lead); writing – original draft (lead); writing – review and editing (equal). **Simon Chamaillé-Jammes:** Conceptualization (equal); formal analysis (supporting); investigation (lead); supervision (lead); funding acquisition (lead); project administration (lead); writing – review and editing (equal).

## Data accessibility statement

Upon publication, data will be made available on a Dryad repository as requested.

## 8 Appendices

### 8.1 Appendix 1. Identification of resting patterns

#### 8.1.1 Identification of the inactive phases

From visual inspection of the activity data, we inferred that, in our study area, the wild boars were nocturnal. The distribution of activity values (ACT) was bimodal for all individuals, with one mode around zero - corresponding to resting - and one mode between 80 and 240 - corresponding to activity (three examples in Figure S1a-b-c). We chose a threshold value (ACT=40) to separate those activity values in two groups such that the least frequent values of activity, i.e. those between 10 and 80, were mostly classified as ‘activity’. We thus kept the ‘resting’ category as constrained as possible. All the individuals considered in the analysis showed a clear nocturnal pattern of activity, with one sequence of low values of activity spanning throughout daytime - sometimes starting a little before sunrise - and one sequence of high values of activity spanning quite precisely from sunset to the start of the next sequence of inactivity, in the morning (three examples in Figure S1d-e-f).

Based on the observed temporal patterns of activity, we considered that each 24-hour cycle could be divided into two continuous phases: the active and the inactive phase. We identified the time boundaries of these phases by applying the same treatment to each animal’s activity data. We first applied a threshold on the raw activity data (one example in Figure S2a) to obtain a binary time series with values being either ‘active’ (ACT>40) or ‘resting’ (ACT≤40) behavior, each value corresponding to a 5min time slot (Figure S2b). Many short sequences of ‘active’ behavior were typically spread across what would be the inactive phase, i.e. during the day, and conversely, some short ‘resting’ sequences existed within what would be the ‘active’ phase, i.e. during the night. We therefore simplified the time series by smoothing out the short sequences that we considered irrelevant for the determination of the time boundaries of ‘active’ and ‘inactive’ phases. We did so by iteratively switching the category (‘resting’ or ‘active’) of the shortest sequence in the time series, until the shortest sequence in the entire time series was 1 hour long (Figure S2c). Five minutes time slots with intermediate activity values (between 10 and 80) were rarely consecutive. For that reason, the outcome of the last processing step was robust to the choice of a different threshold value to identify ‘active’ and ‘resting’ behaviors. Finally, we determined the times of the starting and ending of the ‘inactive’ and ‘active’ phases, so that they matched as closely as possible the pattern observed visually on the raw data (Figure S2d, E and F). We used the following three rules to attribute each sequence of the time series to one or the other phase: (i) All sequences of ‘resting’ behavior overlapping daytime, and (ii) all sequences of ‘active’ behavior spanning entirely within daytime were considered part of the inactive phase. In addition, (iii) whenever a sequence of ‘resting’ behavior restricted to nighttime lasted more than 1.5h, and was separated from the rest of the ‘inactive’ phase by an ‘active’ sequence less than 1.5h long, both the ‘resting’ and the ‘active’ sequences were added to the ‘inactive’ phase. The result of this last step closely matched the pattern expected by visual inspection of the raw data (Figure S2d).

#### 8.1.2 Identification of the resting sites

After having determined the start and end times of the daily inactive phases, we used GPS data to identify where resting had occurred each day. Often, resting occurred at a single resting site’ (RS) throughout the inactive phase (example in Figure S3a). In some instances, however, wild boars did move from an initial RS, to settle in a secondary RS, during the inactive phase, in what we called a ‘relocation’ (example in Figure S3b). Relocations would obviously be associated with important values of ACT. To identify them, we compared the location of the animal before and after the bouts of activity that we considered sufficiently ‘active’ to be considered a ‘possible relocation’ (hereafter PR, see Figure S3c). We considered a PR whenever a continuous ‘active’ bout reached a cumulative ACT value of at least 100. Between PRs, the ACT signal was too low for the animal to have possibly traveled from one RS to another. We compared the mean GPS positions before and after the first PR of each inactive phase. If they were further apart than 100m, we considered that a relocation had occurred. If not, we discarded the PR and applied the same method to the next PR in the inactive phase. Once we had identified all the relocations, we estimated the exact location of the different RSs to be the average of the GPS positions recorded during the inactive phase, between each relocation (Figure S3c).

**Figure S1.**
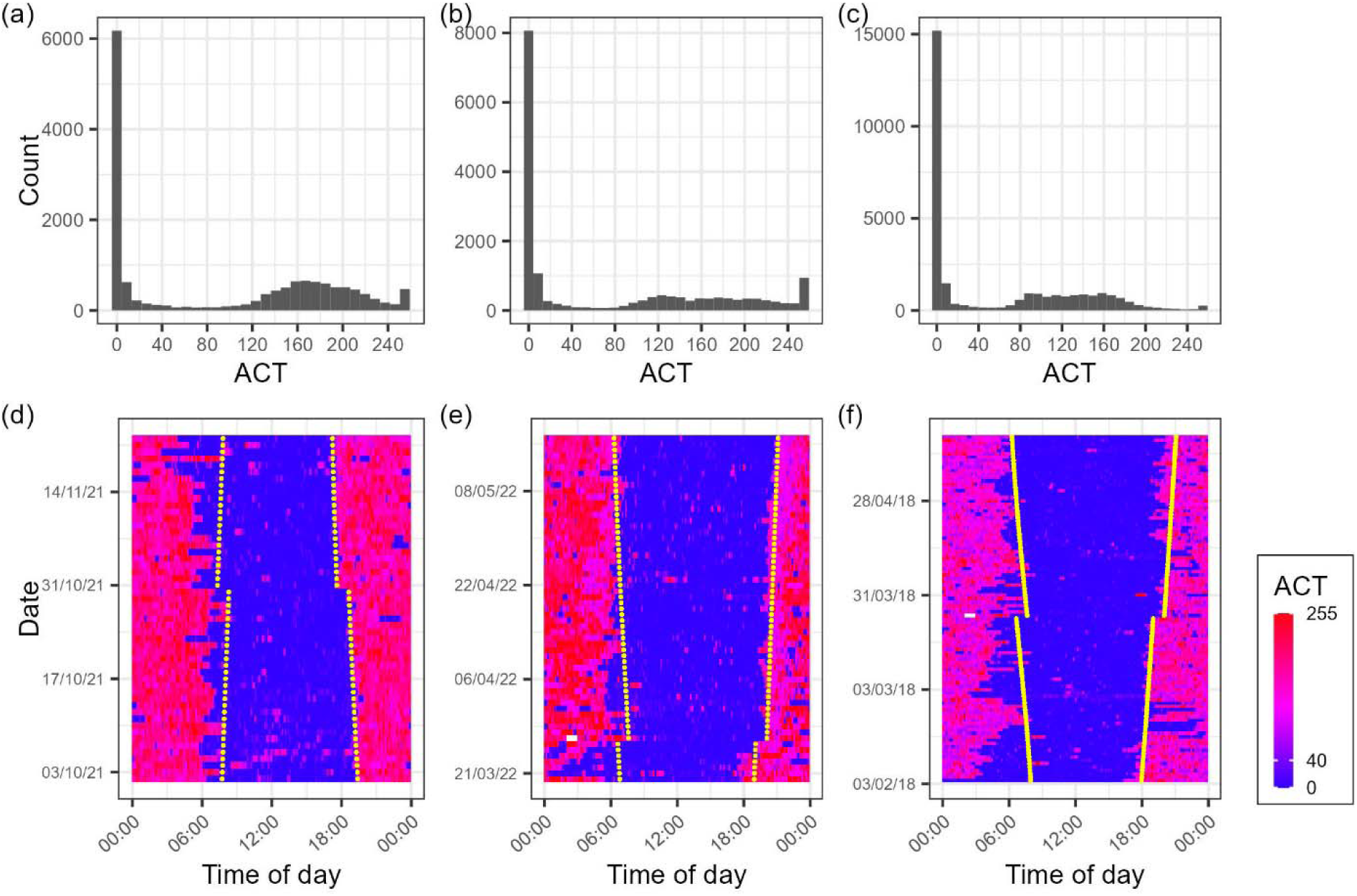
Patterns of activity of three wild boars. (a-b-c) Distribution of the raw ACT values, for three individuals from our study. (d-e-f) Temporal patterns of activity of the animals presented respectively in (a), (b) and (c). Each date (by rows) is divided into 5min slots that are colored according to their level of activity, ranging from blue (ACT = 0) to red (ACT = 255). The yellow dots indicate, for each date, the times of sunrise and sunset. Times are in local time, which includes daylight savings (note the discontinuities in sunrise and sunset times).

**Figure S2.**
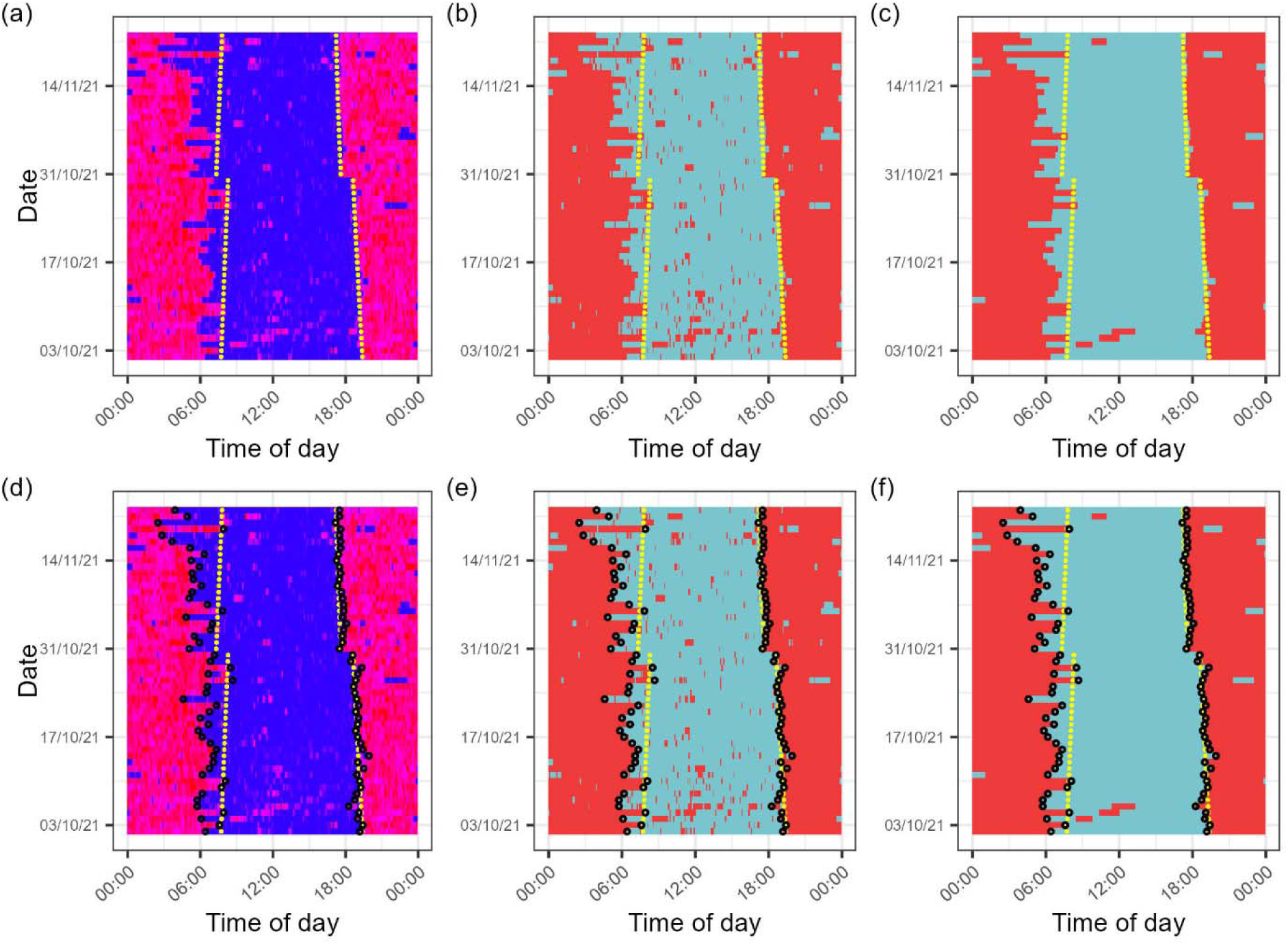
Visual description of the steps taken to determine the timing of the ‘inactive’ phases. (a) Temporal pattern of activity of one wild boar, as raw data. The scale used is the same as in Figure S1. (b) Result of the application of a threshold on the time series of raw data, to distinguish ‘active’ (ACT>40, in red) and ‘inactive’ (ACT<=40, in light blue) behaviors. (c) Result of the smoothing processing step applied to the time series shown in (b). (d), (e) and (f) respectively show how the start and end times of the ‘active’ and ‘inactive’ phases (black circles) fit on the time series presented in (a), (b) and (c). The yellow dots indicate, for each date, the times of sunrise and sunset.

**Figure S3.**
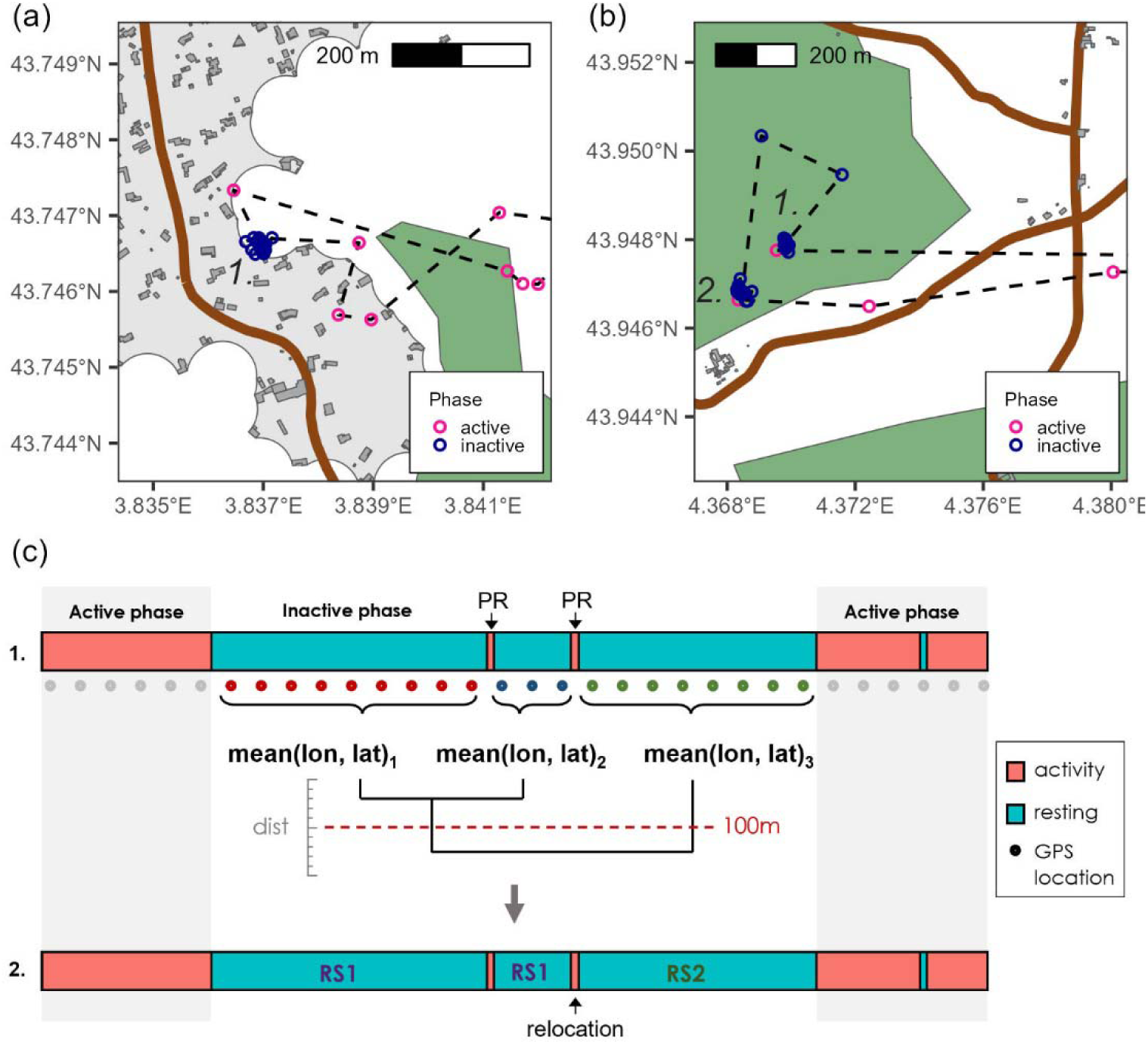
Identification of the relocations. (a) Trajectory of a wild boar (dashed line), including an inactive phase with no relocation, and thus a single resting site (1). (b) Trajectory of a wild boar (dashed line), including an inactive phase with a relocation, in which the animal traveled from a primary resting site (1) to a secondary resting site (2). In (a) and (b), buildings are represented in dark grey, villages in light grey, roads in brown, and densely vegetated patches in dark green. (c) Method of identification of the relocations. We focus on a time series of ‘active’ and ‘resting’ bouts (1). During the inactive phase, between the potential relocations (PR), the distances (dist) between mean GPS positions (mean(lon, lat)) are calculated. Whenever dist exceeds 100m, we consider that the PR corresponds to a relocation. This method enables the identification of the resting sites (RS1 and RS2) used throughout the inactive phase (2).

### 8.2 Appendix 2. Identification of densely vegetated patches

We retrieved spatial information on vegetation cover from CORINE Land Cover 2018 (https://land.copernicus.eu/pan-european/corine-land-cover/clc2018). The following list shows the categories that we included in the definition of ‘densely vegetated patches’, together with the percentage of the land they occupied in our study area:

- Broad-leaved forest (14.5%)
- Coniferous forest (2.4%)
- Mixed forest (4.4%)
- Sclerophyllous vegetation (12.7%)
- Transitional woodland-shrub (4.1%)
- Land principally occupied by agriculture, with significant areas of natural vegetation (3.6%)
- Natural grasslands (2.2%)
- Moors and heathland (0.3%)

Although usually associated with low vegetation, the ‘natural grasslands’ and the ‘moors and heathland’ categories corresponded, in our study area, to areas of regrowth, offering suitable vegetation cover for wild boars to hide. This is why we considered relevant to group them together with taller vegetation layers in this study.

### 8.3 Appendix 3. Sound extracts

All the sound extracts mentioned in the paper are available for listening at the following address: https://doi.org/10.5281/zenodo.7660803. The tracks that were extracted from 40 minutes before to 5 minutes after a relocation was initiated are labelled as ‘relocation’. The tracks labelled as ‘control’ were extracted at the same time of day as the ‘relocation’ tracks, but on different dates, when relocations did not occur. We also make available an additional track, corresponding to a dog barking, with the wild boar not reacting to it. It was identified upon opportunistic listening of the dataset.

## References

Bojarska, K., Maugeri, L., Kuehn, R., Król, W., Theuerkauf, J., Okarma, H. & Gula, R. (2021). Wolves under cover: The importance of human-related factors in resting site selection in a commercial forest. Forest Ecology and Management, 497(119511), doi:https://doi.org/10.1016/j.foreco.2021.119511.

Brivio, F., Grignolio, S., Brogi, R., Benazzi, M., Bertolucci, C. & Apollonio, M. (2017). An analysis of intrinsic and extrinsic factors affecting the activity of a nocturnal species: The wild boar. Mammalian Biology, 84, pp. 73–81, doi:https://doi.org/10.1016/j.mambio.2017.01.007.

Burger, A. L., Fennessy, J., Fennessy, S. & Dierkes, P. W. (2020). Nightly selection of resting sites and group behavior reveal antipredator strategies in giraffe. Ecology and Evolution, 10(6), pp. 2917–2927, doi:https://doi.org/10.1002/ece3.6106.

Campbell, S. S. & Tobler, I. (1984). Animal sleep: A review of sleep duration across phylogeny. Neuroscience & Biobehavioral Reviews, 8(3), pp. 269–300, doi:https://doi.org/10.1016/0149-7634(84)90054-X.

Chamaillé-Jammes, S. (2020). A reformulation of the selection ratio shed light on resource selection functions and leads to a unified framework for habitat selection studies. bioRxiv, doi:https://doi.org/10.1101/565838.

Gaynor, K. M., Hojnowski, C. E., Carter, N. H. & Brashares, J. S. (2018). The influence of human disturbance on wildlife nocturnality. Science, 360(6394), pp. 1232–1235, doi:https://doi.org/10.1126/science.aar7121.

Glass, T. W., Breed, G. A., Robards, M. D., Williams, C. T. & Kielland, K. (2021). Trade-off between predation risk and behavioural thermoregulation drives resting behaviour in a cold-adapted mesocarnivore. Animal Behaviour, 175, pp. 163–174, doi:https://doi.org/10.1016/j.anbehav.2021.02.017.

Ikeda, T., Kuninaga, N., Suzuki, T., Ikushima, S. & Suzuki, M. (2019). Tourist-wild boar (Sus scrofa) interactions in urban wildlife management. Global Ecology and Conservation, 18, p. e00617, doi:https://doi.org/10.1016/j.gecco.2019.e00617.

Johann, F., Handschuh, M., Linderoth, P., Dormann, C. F. & Arnold, J. (2020). Adaptation of wild boar (Sus scrofa) activity in a human-dominated landscape. BMC Ecology, 20(4), doi:https://doi.org/10.1186/s12898-019-0271-7.

Keuling, O., Stier, N. & Roth, M. (2008). How does hunting influence activity and spatial usage in wild boar Sus scrofa L.? European Journal of Wildlife Research, 54(4), pp. 729–737, doi:https://doi.org/10.1007/s10344-008-0204-9.

Latorre, L., Miquel, J. & Chamaillé-Jammes, S. (2021). MEMS based Low-Power Multi-Sensors device for Bio-Logging Applications. In: 2021 Symposium on Design, Test, Integration & Packaging of MEMS and MOEMS (DTIP). Presented at the 2021 Symposium on Design, Test, Integration & Packaging of MEMS and MOEMS (DTIP), pp. 1–4, doi:https://doi.org/10.1109/DTIP54218.2021.9568669.

Lima, S. L., Rattenborg, N. C., Lesku, J. A. & Amlaner, C. J. (2005). Sleeping under the risk of predation. Animal Behaviour, 70(4), pp. 723–736, doi:https://doi.org/10.1016/j.anbehav.2005.01.008.

Lowry, H., Lill, A. & Wong, B. B. M. (2013). Behavioural responses of wildlife to urban environments. Biological Reviews, 88(3), pp. 537–549, doi:https://doi.org/10.1111/brv.12012.

Lutermann, H., Verburgt, L. & Rendigs, A. (2010). Resting and nesting in a small mammal: sleeping sites as a limiting resource for female grey mouse lemurs. Animal Behaviour, 79(6), pp. 1211–1219, doi:https://doi.org/10.1016/j.anbehav.2010.02.017.

Maillard, D. & Fournier, P. (1995). Effects of shooting with hounds on size of resting range of Wild boar (Sus scrofa L.) groups in mediterranean habitat. IBEX Journal of Mountain Ecology, 3(102), p. e107.

Markham, A. C., Alberts, S. C. & Altmann, J. (2016). Haven for the night: sleeping site selection in a wild primate. Behavioral Ecology, 27(1), pp. 29–35, doi:https://doi.org/10.1093/beheco/arv118.

Marzano, M. & Dandy, N. (2012). Recreationist behaviour in forests and the disturbance of wildlife. Biodiversity and Conservation, 21(11), pp. 2967–2986, doi:https://doi.org/10.1007/s10531-012-0350-y.

Mortlock, E., Silovský, V., Güldenpfennig, J., Faltusová, M., Olejarz, A., Börger, L., Ježek, M., Jennings, D. J. & Capellini, I. (2022). Individual identity and environmental conditions explain different aspects of sleep behaviour in wild boar. bioRxiv, doi:https://doi.org/10.1101/2022.11.23.517569.

Ohashi, H., Saito, M., Horie, R., Tsunoda, H., Noba, H., Ishii, H., Kuwabara, T., Hiroshige, Y., Koike, S., Hoshino, Y., Toda, H. & Kaji, K. (2014). Erratum to: Differences in the activity pattern of the wild boar Sus scrofa related to human disturbance. European Journal of Wildlife Research, 60(567), doi:https://doi.org/10.1007/s10344-012-0661-z.

Podgórski, T., Baś, G., Jędrzejewska, B., Sönnichsen, L., Śnieżko, S., Jędrzejewski, W. & Okarma, H. (2013). Spatiotemporal behavioral plasticity of wild boar (Sus scrofa) under contrasting conditions of human pressure: primeval forest and metropolitan area. Journal of Mammalogy, 94(1), pp. 109–119, doi:https://doi.org/10.1644/12-MAMM-A-038.1.

Riede, S. J., van der Vinne, V. & Hut, R. A. (2017). The flexible clock: predictive and reactive homeostasis, energy balance and the circadian regulation of sleep–wake timing. Journal of Experimental Biology, 220(5), pp. 738–749, doi:https://doi.org/10.1242/jeb.130757.

Rosalino, L. M., Teixeira, D., Camarinha, C., Pereira, G., Magalhães, A., Castro, G., Lima, C. & Fonseca, C. (2022). Even generalist and resilient species are affected by anthropic disturbance: evidence from wild boar activity patterns in a Mediterranean landscape. Mammal Research, 67(3), pp. 317–325, doi:https://doi.org/10.1007/s13364-022-00632-8.

Saïd, S., Tolon, V., Brandt, S. & Baubet, E. (2012). Sex effect on habitat selection in response to hunting disturbance: the study of wild boar. European Journal of Wildlife Research, 58(1), pp. 107–115, doi:https://doi.org/10.1007/s10344-011-0548-4.

Schmidt, M. H. (2014). The energy allocation function of sleep: A unifying theory of sleep, torpor, and continuous wakefulness. Neuroscience & Biobehavioral Reviews, 47, pp. 122–153, doi:https://doi.org/10.1016/j.neubiorev.2014.08.001.

Scillitani, L., Monaco, A. & Toso, S. (2010). Do intensive drive hunts affect wild boar (Sus scrofa) spatial behaviour in Italy? Some evidences and management implications. European Journal of Wildlife Research, 56(3), pp. 307–318, doi:https://doi.org/10.1007/s10344-009-0314-z.

Siegel, J. M. (2009). Sleep viewed as a state of adaptive inactivity. Nature Reviews Neuroscience, 10(10), pp. 747–753, doi:https://doi.org/10.1038/nrn2697.

Sodeikat, G. & Pohlmeyer, K. (2007). Impact of drive hunts on daytime resting site areas of wild boar family groups (Sus scrofa L.). Wildlife Biology in Practice, 3, pp. 28–38, doi:https://doi.org/10.2461/wbp.2007.3.4.

Stillfried, M., Gras, P., Börner, K., Göritz, F., Painer, J., Röllig, K., Wenzler, M., Hofer, H., Ortmann, S. & Kramer-Schadt, S. (2017). Secrets of Success in a Landscape of Fear: Urban Wild Boar Adjust Risk Perception and Tolerate Disturbance. Frontiers in Ecology and Evolution, 5(157), doi:https://doi.org/10.3389/fevo.2017.00157.

Thurfjell, H., Spong, G. & Ericsson, G. (2013). Effects of hunting on wild boar Sus scrofa behaviour. Wildlife Biology, 19(1), pp. 87–93, doi:https://doi.org/10.2981/12-027.

Tolon, V., Dray, S., Loison, A., Zeileis, A., Fischer, C. & Baubet, E. (2009). Responding to spatial and temporal variations in predation risk: space use of a game species in a changing landscape of fear. Canadian Journal of Zoology, 87(12), pp. 1129–1137, doi:https://doi.org/10.1139/Z09-101.

Wam, H. K., Eldegard, K. & Hjeljord, O. (2012). From overlooking to concealed: predator avoidance in an apex carnivore. European Journal of Wildlife Research, 58(6), pp. 1001–1003, doi:https://doi.org/10.1007/s10344-012-0670-y.

Wilson, M. W., Ridlon, A. D., Gaynor, K. M., Gaines, S. D., Stier, A. C. & Halpern, B. S. (2020). Ecological impacts of human-induced animal behaviour change. Ecology Letters, 23(10), pp. 1522–1536, doi:https://doi.org/10.1111/ele.13571.

Wittemyer, G., Keating, L. M., Vollrath, F. & Douglas-Hamilton, I. (2017). Graph theory illustrates spatial and temporal features that structure elephant rest locations and reflect risk perception. Ecography, 40(5), pp. 598–605, doi:https://doi.org/10.1111/ecog.02379.

